# Co-occurrence Networks and Topological Analyses Revealed Microbiome Structures in Soybean Roots from a Louisiana Soybean Field Severely Damaged by Prolonged High Temperatures and Drought Stress

**DOI:** 10.1101/2024.01.16.575963

**Authors:** Sandeep Gouli, Aqsa Majeed, Jinbao Liu, David Moseley, M. Shahid Mukhtar, Jong Hyun Ham

**Author notes:** Corresponding authors /. ^§^ Equally contributed.

## Abstract

Drought stress has a significant impact on agricultural productivity, affecting key crops such as soybeans, the second most widely cultivated crop in the United States. We conducted endophytic and rhizospheric microbial diversity analyses in soybean plants cultivated during the 2023 growing season, amid extreme weather conditions of prolonged high temperatures and drought in Louisiana. Specifically, we collected surviving and non-surviving soybean plants from two plots of a Louisiana soybean field severely damaged from the extreme heat and drought condition in 2023. We did not observe any significant difference in the microbial diversity of rhizosphere between surviving and non-surviving plants. However, we found obvious differences in the structure of endophytic microbial community in root tissues between the two plant conditions. Especially, the bacterial genera of Proteobacteria, *Pseudomonas* and *Pantoea*, were predominant in the surviving root tissues, while the bacterial genus *Streptomyces* was conspicuously dominant in the non-surviving (dead) root tissues. Co-occurrence patterns and network centrality analyses enabled us to discern the intricate characteristics of operational taxonomic units (OTUs) within endophytic and rhizospheric networks. Overall, this study advances our understanding of the intricate relationship between bacteria and plants under drought stress, paving the way for future research to investigate the importance of microbial diversity in drought affected regions such as Louisiana.

## Introduction

Drought stress is one of the most important environmental factors that significantly impact agricultural productivity, leading to a potential yield loss of 50% yield loss [1, 2]. This can have severe implications for global feed security. The second most widely cultivated crop in the United States is soybeans, representing 32% of the overall cultivated land. However, under drought conditions, soybeans can experience a reduction in yield of up to 100% [1]. As the crisis of global climate change continues to escalate, the frequency and severity of drought events are intensifying, presenting a greater threat to crop yields and overall agricultural sustainability. Consequently, understanding the mechanisms underlying plant responses to drought stress is imperative for developing resilient soybean varieties and implementing effective management strategies. Utilizing beneficial microbes has emerged as a promising approach to enhance crop resilience against environmental stress, including drought [3].

Assessing the microbial diversity and population of both endophytes and the rhizosphere in crop plants across various environments is crucial to harnessing their potential use in enhancing plant growth and resilience against both biotic and abiotic stress [3]. The progress in high-throughput sequencing and bioinformatics tools allows for the evaluation of operational taxonomic units (OTU), amplicon sequence variants (ASV), or species, along with their respective abundances [4]. Leveraging correlation-based and graphical models, among other methods, co-occurrence network analysis is applied to illustrate microbial relationships within diverse spatiotemporal niches [5]. Meanwhile, network topological features i.e. modularity and connectivity, and several of such network parameters can serve as indicators of significant nodes in the network [6]. These encompass degree, indicating the number of connections a node possesses; betweenness, representing the fraction of the shortest paths passing through a node, among other centrality measures [7]. Recently, weight *k*-shell decomposition network analysis was shown to be more effective in discovering fast information-spreading nodes [8].

During the 2023 growing season, most regions of Louisiana experienced extreme weather conditions of high temperature and drought for an unprecedentedly extended period. From these abnormally hot and dry weather conditions, some of the soybean fields at the Dean Lee Research Station of the LSU AgCenter in Alexandria, Louisiana, exhibited devastating damages and most of the plants could not survive, causing a more than 90% yield loss (Fig. 1). The primary objective of this study is to investigate the microbiome structure of soybean roots under natural drought and heat stresses in the field, particularly during this year’s severe environmental growing season. As there were no healthy plants at a similar growth stage cultivated in nearby well-irrigated plots, soybean plants were collected from two damaged plots only. However, both surviving and non-surviving plants were sampled from each plot for comparative analysis.

This report describes the prokaryotic microbiome structures associated with the soybean roots that survived the natural heat and drought stresses in the field condition of central Louisiana during the 2023 growing season, compared with those from non-surviving plants in the same plot. We present here the alpha and beta diversity and the co-occurrence network of the microbiome structures based on the 16S ribosomal DNA (16S rDNA) sequence data. We also discuss here the main characteristics of the microbiome associated with the hot and dry field conditions.

**Figure 1.**
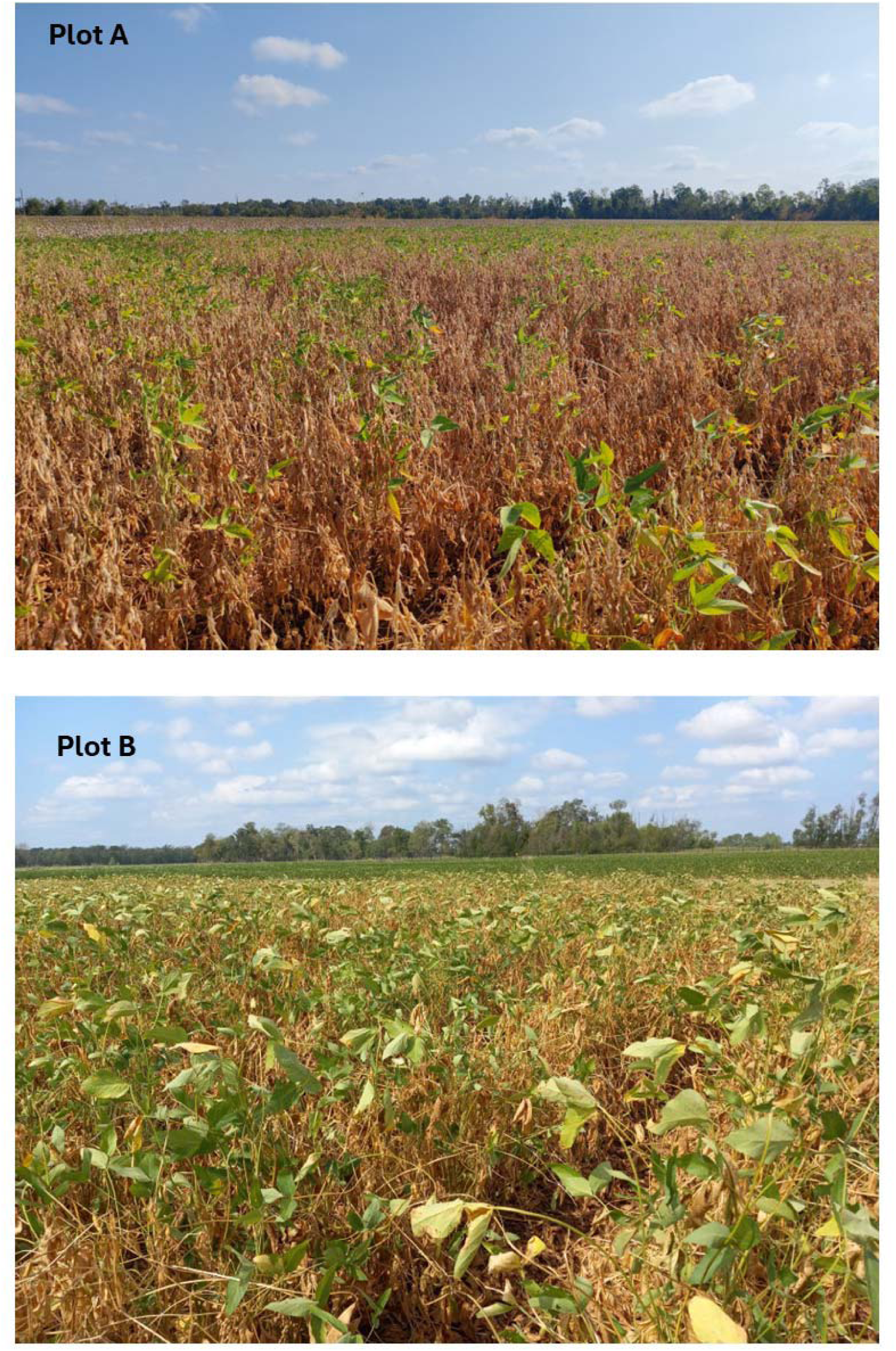
The drought-damaged soybean field where the plant samples were collected (Alexandria, Louisiana). Plot A and Plot B were neighboring each other.

## Materials and Methods

### Plant Sample Collection and Preservation

Soybean plant samples were collected from two neighboring plots (Plot A and Plot B) at the Dean Lee Research Station, Alexandria, Louisiana (31.11°N, 92.24°W), which were severely damaged from the extended period of high temperatures and dry weather conditions during the 2023 growing season (Figures 1 and 2). Five surviving and five dead plants were collected at the R6-R7 stage (∼ 4 months after planting) from both Plot A and Plot B, totaling ten plant samples per plot. Plot A experienced severe drought stress, resulting in the survival of only a few green plants, while Plot B exhibited less severe drought stress with more green plants than Plot A (Figure 1). Soil types in both plots included Latanier silty clay loam at the front and Moreland clay at the back. Tillage was conducted in March, and the planting was done on May 4, 2023, for both plots. Different crop varieties were cultivated in each plot: AG49XF3 (Bayer CropScience) for Plot A and P5554RX (Progeny Ag) for Plot B. The collected plant and soil samples were carefully encased in large plastic bags and transported to the laboratory, where they were temporarily stored in a cold room at 4^0^C before processing.

**Figure 2.**
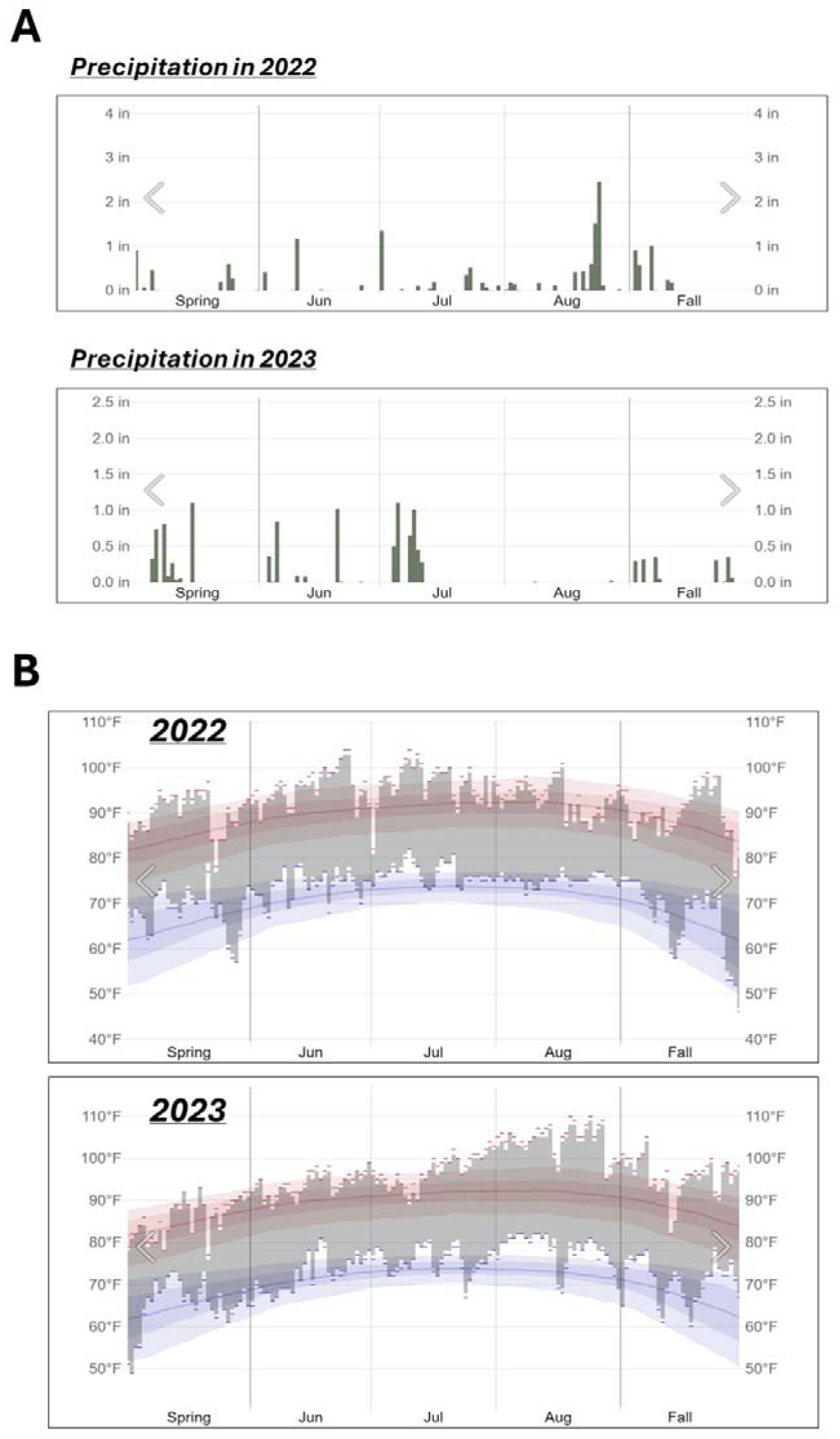
The records of the rainfalls (A) and temperatures (B) in Alexandria, Louisiana, during the 2023 growing season. These graphs were obtained from Weather Spark (https://weatherspark.com).

### Field Management

#### Insecticide and herbicide applications

In both plots, insecticides were applied on two dates: Moccasin MTZ (United Phosphorous, PA, USA) on July 17, 2023, at a rate of 0.41 ml/m^2^, and Endigo (Syngenta, DE, USA) on July 24, 2023, at a rate of 0.033 ml/m^2^. Herbicide applications were administered on May 8, 2023, with Varsity (Innvictis Crop Care, Idaho, USA) at a rate of 0.015 ml/m^2^, Derive (Innvictis Crop Care, Idaho, USA) at a rate of 0.045 ml/m^2^, Zidua (BASF Agricultural Solutions, LA, USA) at a rate of 0.018 ml/m^2^, and Fever (Innvictis Crop Care, Idaho, USA) at a rate of 0.234 ml/m^2^. Charger Max (WinField United, MN, USA) was applied at a rate of 0.146 ml/m^2^ on May 9, 2023, and on June 15, 2023, Sentris (BASF Agricultural Solutions, LA, USA) at a rate of 0.06 ml/m^2^, Engenia (BASF Agricultural Solutions, LA, USA) at a rate of 0.09 ml/m^2^, and Roundup Powermax (Bayer Crop Science, MO, USA) at a rate of 0.23 ml/m^2^ were administered.

#### Fungicide and fertilizer applications

On June 30, 2023, a fungicide, Stratego Yield (Bayer CropScience, MO, USA), was applied at a rate of 0.034 ml/m^2^ grams in both fields, supplemented with a 0.25% nonionic surfactant. The fertilizer used in both fields was a composition of N-P-K at 0-18-36.

### DNA sample preparation

For DNA extraction from the rhizospheric soil, the soil closely attached to the plant roots was collected in 50 ml falcon tubes using a sterile spatula and gloves to prevent contamination. Each sample was labeled appropriately according to the status of each plant sample. 0.25 g of the collected soil sample was used for DNA extraction. The DNA extraction was performed using DNeasy PowerSoil Pro Kit (Qiagen GmbH, Hilden, Germany) following the manufacturer’s instructions. After DNA extraction, the quantity and quality of each DNA sample were assessed using a NanoDrop 1000 Spectrophotometer (Thermo Scientific, Wilmington, DE, USA). For DNA extraction from the inner part of plant roots (endosphere), the plant root samples that remained after collection of the rhizospheric soil were surface sterilized by immersing them in 70% ethanol and subsequently rinsing with double-distilled water to ensure the removal of microbes on the surface of root tissue. Under aseptic conditions and using sterile scissors the surface-sterilized roots were cut into pieces (∼ 1 to 2 mm thick) to access the inner endospheric part. DNA extraction of the plant tissue samples for the endosphere was conducted utilizing a DNeasy Plant Mini Kit (Qiagen GmbH, Hilden, Germany) following the manufacturer’s instructions. Subsequently, the DNA samples extracted from the endosphere samples were quantified using a NanoDrop 1000 Spectrophotometer (Thermo Scientific, Wilmington, DE, USA). Extracted DNA was then sequenced for 16S rDNA sequencing.

### Library Construction and high-throughput DNA sequencing of 16S rDNA

The V4 regions of bacterial 16S rDNA were amplified using the primers 515F (5’-GTGYCAGCMGCCGCGGTAA-3’) and 806R (5’-GGACTACNVGGGTWTCTAAT-3’) designed in published protocols [9, 10]. Thirty nanograms of isolated DNA were used as the template for the PCR amplification reaction and subsequent library construction. The resultant products were purified using Agencourt AMPure XP beads and subsequently measured using an Aligent 2100 bioanalyzer to determine their size and concentration. Samples that passed the quality control were sequenced on DNBSEQ-G99 platform (MGI Tech, ShenZhen, China) using a 300bp paired-end strategy.

### Raw Data Import, Quality checking, and ASV feature table Construction

Raw paired-end reads (FASTQ) from the original DNA fragments were imported in Qiime2 v 2023.2 software [11]. Paired-end reads for all 20 samples for the dataset Endosphere (root) and 20 samples for Rhizosphere (soil) were imported using the manifest file method. Quality filtering, denoising, and chimeric sequence removal were done using the DADA2 denoising method. To remove low-quality regions of the sequences --**p-trunc-len** parameter was used to truncate each sequence at position 253 in forward and reverse reads. This DADA2 pipeline generated a FeatureTable [Frequency], which contains counts (frequencies) of each unique sequence in each sample in the dataset, and a FeatureData[Sequence], which maps feature identifiers in the Feature Table to these sequences.

### Taxonomy Assignment

To explore the taxonomic composition of the samples, a pre-trained Naïve Bayes classifier and q2-feature classifier plugin were used to assign likely taxonomies to the sequences. This classifier, downloaded from the qiime2 data resources page, was trained on the SILVA OTUs [12] from the V4 (515F/806R) region of sequences. Taxa bar plots were generated using an R package microViz (V. 0.12.0) [13] to visualize the taxonomic composition of each sample and group at Phylum, Family, and Genus classification levels. Bar plots are used to visualize OTUs’ relative abundance. All nonbacterial OTUs’ sequences are filtered out using the feature-table-filtering method in Qiime2.

### Diversity Analysis

To assess the alpha diversity, three different metrics are calculated: “Evenness” estimates the species abundance; “Observed OTU” estimates the number of unique OTUs found in each sample, and “Shannon index” accounts for both richness and evenness. Shannon index value ranges from 0 to 1. Lower values indicate high diversity and higher index values do lower diversity. These alpha diversity metrics were calculated using the phyloseq function ‘estimate_richness’ [14] and to visualize diversity results, Boxplots are generated using “qiime2R” [15] and “ggplot2” [16] libraries in R. For beta diversity analysis, Principal Co-ordinates Analysis (PCoA) was performed. PCoA is an unconstrained method, but it does require a distance matrix. In an ecological context, a distance (or more generally a “dissimilarity”) measure indicates how different a pair of (microbial) ecosystems are. This can be calculated in many ways. For this study, Weighted Unifrac distance, Unweighted Unifrac distance, and Generalized UniFrac, “gunifrac were selected to generate PCoA curve to measure the dissimilarity coefficient between pairwise samples, which are phylogenetic measures used extensively in recent microbial community sequencing projects. An R package microViz (V. 0.12.0) [13] was used to generate these plots. The <u>UniFrac family</u> of methods was employed to determine dissimilarities. The approach considers the phylogenetic relatedness of taxa/sequences in samples. Conversely, un-weighted UniFrac, dist_calc(dist = “unifrac”) disregards the relative abundance of taxa, and highlights solely on their presence (detection) or absence. This renders it particularly sensitive to rare taxa, sequencing artifacts, and abundance filtering choices. For assessing dissimilarities, on the other hand, weighted UniFrac, denoted as “wunifrac,” places (potentially excessive) emphasis on highly abundant taxa. The Generalized UniFrac, labeled as “gunifrac,” achieves a balance between the extremes of unweighted and weighted UniFrac.

### Co-occurrence microbial network analysis

An R package called ggClusterNet [17] was used to make networks for Endosphere (root) and Rhizosphere (soil) datasets. An integrated function network.2() of ggClusterNet was used for microbial network data mining and visualization. Briefly, to calculate network correlation the Spearman method was used, 0.9 correlation and 0.05 P-value thresholds were used to filter the microorganism table. The model_maptree2 layout has been used for the microbial network. Two types of networks were calculated:1) Global network, including all OTU from all samples; 2) Individual network for each of four groups from Endosphere (root) and Rhizosphere (soil) datasets.

## Results and Discussion

### Alpha and beta diversities

We analyzed the alpha and beta diversities of both rhizospheric and endospheric bacterial communities. As we did not find any significant differences in beta diversity among the rhizosphere samples regardless of their surviving status and plot condition (data not shown), our analysis of bacterial community shifted to the root endosphere. Here, we observed significant differences in beta diversity between drought-surviving and non-surviving soybean root tissues in both Plot A and Plot B (p-value = 0.001, permanova, pseudo-F test statistic). Pairwise permanova results further confirmed these distinctions. Notably, alpha diversity analysis revealed evenness within the groups. A closer scrutinizing of taxa bar plots identified seven genera— *Pseudomonas, Pantoea, Streptomyces, Micrococcaceae, Luteimonas, Variovorax,* and *Bacillus*— as being permanently abundant in either surviving or dry/dead plants from both fields (Figures 3 and 4). These findings led us to hypothesize that these bacterial genera might play a crucial role in enhancing soybean tolerance to severe drought stress. A comprehensive bar graph visually illustrates the abundance of these bacterial genera in surviving plants except for *Streptomyces*, which is abundant in dry/dead plants in both fields (Figure 4).

**Figure 3.**
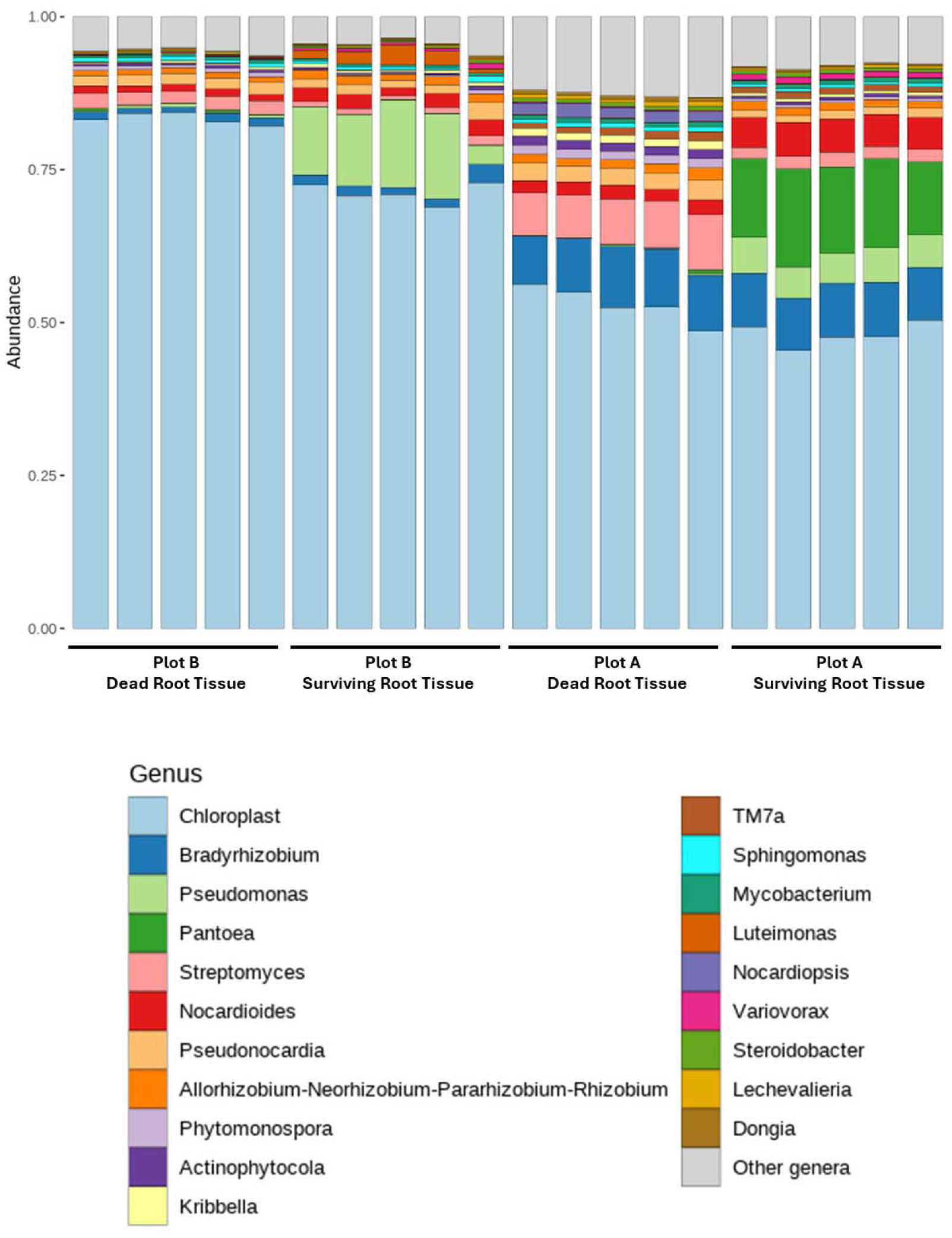
Taxa bar plot showing seven abundant bacterial genera either in surviving or dry/dead soybean root tissues (root endosphere).

**Figure 4.**
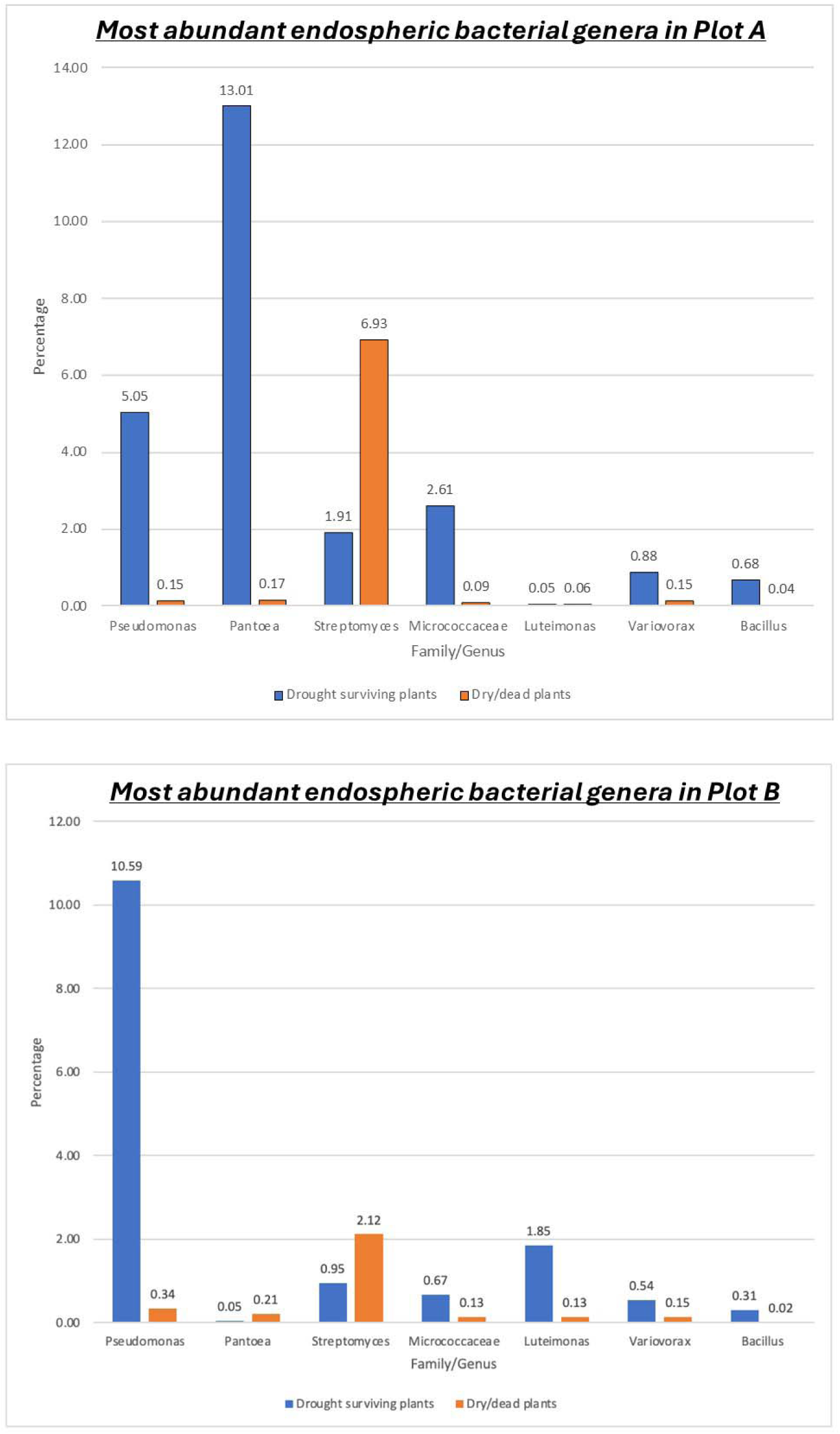
Relative abundance of bacterial genera dominant in the root endosphere of drought-damaged soybean plants.

Among the seven major bacterial genera, Gram-negative bacteria *Pseudomonas* and *Pantoea* were conspicuously predominant in surviving roots, although *Pantoea* showed reduced presence in Plot B, which has less stress damage (Figure 4). In contrast, *Streptomyces* was the genus that was most conspicuously dominant over other bacterial genera (Figure 4).

The predominance of *Pseudomonas* strains in our microbiome analysis is noteworthy given their potential contributions to enhancing soybean tolerance to severe drought stress. Extensive studies have highlighted the ability of *Pseudomonas* strains to produce Volatile Organic Compounds (VOCs) that directly assist plants in withstanding drought and high salinity [18]. Furthermore, *Pseudomonas* strains, such as *P. simiae* AU, have been associated with induced systemic tolerance (IST) in plants against multiple abiotic stresses, aiding in the accumulation of proline and reducing sodium content in roots to cope with osmotic and ionic stress [19]. The formation of biofilms by plant-beneficial *Pseudomonas* spp. has also been identified as a mechanism that improves tolerance to various stresses, including osmotic and oxidative stress, while enabling the production of beneficial secondary metabolites [20, 21]. Additionally, *Pseudomonas fluorescens* DR397, isolated from drought-prone rhizospheric soil, exhibited high metabolic activity under drought conditions and upregulated the expression of genes related to plant growth promotion, resulting in increased shoot and root growth in legume cultivars under drought conditions [22]. Moreover, the application of *Pseudomonas putida* H-2-3 reprogrammed chlorophyll, stress hormones, and antioxidants in abiotic stress-affected soybean plants, improving their growth under saline and drought conditions [23]. Collectively, these findings underlie the crucial role of *Pseudomonas* strains in mitigating drought stress in soybean through various mechanisms, including VOC production, biofilm formation, and plant physiological modulation.

The other predominant bacterial genus *Pantoea* in the surviving root samples of Plot A is also reminiscent of previous studies showing the biological roles of *Pantoea* spp. in protecting plants from abiotic stresses. The colonization of wheat plants by *Pantoea agglomerans,* known for its exopolysaccharide (EPS) production, positively influenced rhizosphere soil aggregation by increasing the RAS/RT ratio and enhancing the water stability of adhering soil aggregates [24]. In a separate study, *Pantoea* strain LTYR-11ZT, isolated from the leaves of the drought-tolerant plant *Alhagi sparsifolia*, exhibited multiple plant growth-promoting (PGP) traits, improving wheat performance under drought conditions. This strain enhanced soluble sugar accumulation, reduced proline and malondialdehyde levels, and decreased chlorophyll degradation in leaves [25]. Moreover, EPS derived from *Pantoea alhagi* NX-11 demonstrated significant improvements in drought resistance in rice seedlings, increasing fresh weight, and relative water content, and enhancing various physiological parameters, including total chlorophyll, proline, and soluble sugar content [26]. Additionally, *Pantoea sp.* YSD J2, isolated from the leaves of *Cyperus esculentus* L. var. *sativus*, exhibited notable plant growth-promoting characteristics, including indole acetic acid production, siderophores generation, and the ability to solubilize phosphate and potassium [27]. These collective findings highlight the diverse roles of *Pantoea* species in promoting plant growth, enhancing drought tolerance, and contributing to sustainable agriculture through various mechanisms, including EPS production and PGP traits.

Our observation that the genus *Streptomyces* was solely prevailing over other major bacterial genera is not surprising, regarding that this bacterial genus is one of the most abundant microorganisms in soil [28, 29]. In dead root tissues, all the bacterial organisms that are dependent on the interactions with living plant tissues would be rapidly replaced with the dominant soil saprophytes, such as *Streptomyces* spp. Our results suggest a preference for the saprophytic lifestyle of *Streptomyces* in colonizing dead or dry plants, contributing to the intriguing dynamics of microbial interactions in stressed plant environments. Further investigations into the specific mechanisms underlying *Streptomyces* preferences and its potential impact on plant health in stressed conditions would be valuable for a comprehensive understanding of its ecological role.

Regarding the other 4 major bacterial genera identified in this study, several previous studies reported their relatedness with plant drought stress, supporting the idea that these bacterial organisms can be good candidate biological materials to augment the resilience of crops to drought stress. Arun et al. [30] reported the isolation of *Micrococcus* spp. from various environmental sources, including soil and plant samples, and highlighted the plant growth-promoting properties of *Micrococcus luteus*. Notably, the strain K39 of *Micrococcus luteus*, isolated from the roots of *Cyperus conglomeratus* in a desert environment, exhibited characteristics relevant to drought stress survival [31]. This finding aligns with our results, where Plot A, experiencing severe drought stress, showed a higher abundance of bacteria from the Micrococcaceae family compared to Plot B. Related to another major genus *Lutemonas*, *L. deserti* sp. nov. is an intriguing species isolated from the desert soil of northern PR China, providing evidence for the presence of *Luteimonas* in the endosphere of drought-stressed plants [32]. This discovery aligns with our observations, suggesting a potential association between *Luteimonas* and plants experiencing drought stress.

*Variovorax paradoxus* strain 5 C-2, characterized by the presence of the enzyme 1-aminocyclopropane-1-carboxylate (ACC) Deaminase, has been associated with notable benefits for plant growth under drought conditions. Belimov et al. [33] reported that this strain contributes to a reduction in ethylene (ET) production, leading to increased nodulation, elevated seed nitrogen content, enhanced xylem abscisic acid (ABA) concentration, improved water content, and ultimately a higher pea yield. The findings suggest that the enzymatic activity of ACC Deaminase in *Variovorax paradoxus* plays a crucial role in modulating hormone signaling both locally in the rhizosphere and systemically within the plant. These observations underscore the potential of *Variovorax paradoxus* as a beneficial rhizobacterium in promoting plant growth and yield, particularly in conditions of soil moisture limitation.

*Bacillus*, particularly *Bacillus thuringiensis* (UFGS2), has demonstrated its ability to mitigate the impacts of drought stress in plants. In soybean, UFGS2-treated plants exhibited higher stomatal conductance and transpiration compared to the control group following drought stress [34]. This suggests a positive influence of *Bacillus thuringiensis* on plant water regulation under water scarcity conditions. Similarly, the combined application of *Pseudomonas putida* and *Bacillus amyloliquefaciens* alleviated drought stress in chickpeas by exhibiting multiple plant growth-promoting traits, including ACC deaminase activity, mineral solubilization, hormone production, biofilm formation, and siderophore production [35]. Moreover, *Bacillus paralicheniformis* strain FMCH001 demonstrated the potential to enhance water use efficiency, nutrient uptake, root growth, photosynthesis rate, C:N ratio, and overall plant-water relations in soybean, making it a promising candidate for sustaining plant growth in water-limited conditions [36]. Additionally, *Bacillus pumilus* strain SH-9, identified as a drought-tolerant variant, positively influenced soybean growth under drought stress by modulating the expression of phytohormone genes and antioxidant profiles [37]. Furthermore, *Bacillus subtilis* emerged as a beneficial bacterium promoting the growth of common beans and maize while increasing water use efficiency. This bacterium enhanced leaf water content, regulated stomatal activity, and decreased antioxidant activities without compromising photosynthetic rates [38]. In summary, *Bacillus* species, including *Bacillus thuringiensis*, *Bacillus amyloliquefaciens*, *Bacillus paralicheniformis*, *Bacillus pumilus*, and *Bacillus subtilis*, exhibit promising capabilities in alleviating drought stress and enhancing plant growth under challenging environmental conditions. These findings emphasize the potential of *Bacillus*-based strategies for sustainable agriculture in water-limited regions.

### Co-occurrence networks analyses

Network analyses play crucial roles in deciphering co-occurrence patterns across microbial taxa within complex communities. They also facilitate determining positive and negative interactions among diverse taxa [17]. Here, we constructed two sets of co-occurrence networks: endophytic and rhizospheric networks (Figure 5). Within each category, four networks were developed, namely, surviving plot A, non-surviving plot A, surviving plot B, and non-surviving plot B. In total, eight co-occurrence networks were constructed using the significant correlations among phyla (Spearman’s correlation coefficient *r*lJ>lJ0.9, *p*lJ<lJ0.05). Each network exhibited varied network features (Figure 5, Table 1). In general, rhizospheric networks consist of a greater number of nodes, edges, and clusters compared to all endophytic networks, with the highest numbers observed in surviving plot A. Conversely, among endophytic networks, non-surviving plot A displayed the highest number of nodes, edges, and clusters (Figure 5). These findings suggest the existence of varied levels of OTU complexity in endophytic and rhizospheric networks in plot A. These network features are consistent with the average degree of each network. Intriguingly, non-surviving plot A of endophytic networks showed the highest numbers of negative edges, indicating the existence of both synergistic and antagonistic relations among diverse OTUs. The edge density, a network characteristic that represents the proportion of possible relationships in the network, signifies the intricacy of the network and the presence of robust interactions among diverse OTUs. Among the eight networks studied, surviving plot B of endophytic networks exhibited the highest edge density of 0.07, while the lowest value of 0.04 was observed in non-surviving plot A of endophytic networks. The highest modularity, a network feature assessing the network’s structure, was identified in surviving plot A (12.04) of rhizospheric networks. Conversely, the lowest modularity was observed in surviving plot B of endophytic networks (Figure 5 and Table 1). In summary, the examination of co-occurrence patterns and network centrality analyses allowed us to recognize the intricate nature of operational taxonomic units (OTUs) within each network category.

**Figure 5.**
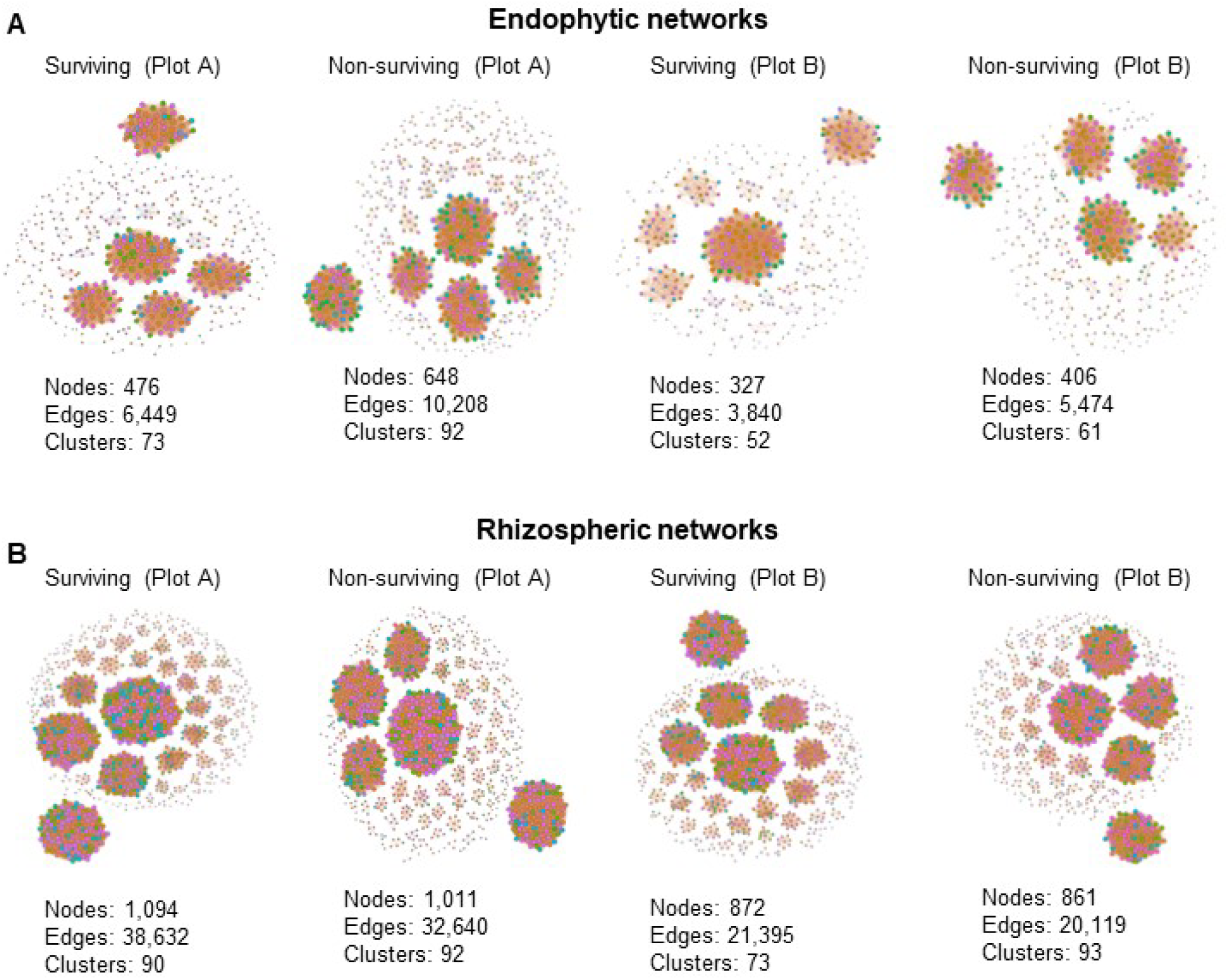
The co-occurrence networks analyses. Four groups each for endophytic (A) and rhizospheric networks (B) are illustrated. Individual types of networks within each category are indicated. Nodes, edges, and clusters for each network are specified. The color of nodes signifies OTUs from the same module in each network, while line color indicates positive (orange) and negative (blue) correlation coefficients. Network construction employed Spearman’s correlation coefficient, with RlJ>lJ0.9 and plJ<lJ0.05 as criteria.

**Table 1.** Network features of diverse co-occurrence networks.

|  | Endophytic networks |  |  |  | Rhizospheric networks |  |  |  |
| --- | --- | --- | --- | --- | --- | --- | --- | --- |
| Centrality Features | Surviving<br>(Plot A) | Non-surviving<br>(Plot A) | Surviving<br>(Plot B) | Non-surviving<br>(Plot B) | Surviving<br>(Plot A) | Non-surviving<br>(Plot A) | Surviving<br>(Plot B) | Non-surviving<br>(Plot B) |
| Nodes | 476 | 648 | 327 | 406 | 1094 | 1011 | 872 | 861 |
| Edges | 6449 | 10208 | 3840 | 5474 | 38632 | 32640 | 21395 | 20119 |
| Positive Edges | 6320 | 10050 | 3765 | 5426 | 38590 | 32543 | 21352 | 19992 |
| Negative Edges | 129 | 158 | 75 | 48 | 42 | 97 | 43 | 127 |
| Number of Clusters | 73 | 92 | 52 | 61 | 90 | 92 | 73 | 93 |
| Connectance (Edge Density) | 0.05704556 | 0.048695785 | 0.07204368 | 0.06658152 | 0.06461595 | 0.063930429 | 0.0563388 | 0.054341896 |
| Average degree (Average K) | 27.0966387 | 31.50617284 | 23.4862385 | 26.9655172 | 70.6252285 | 64.56973294 | 49.071101 | 46.7340302 |
| Average Path Length | 1 | 1 | 1 | 1 | 1 | 1 | 1 | 1 |
| Diameter | 1 | 1 | 1 | 1 | 1 | 1 | 1 | 1 |
| Mean Clustering Coefficient (Average CC) | 1 | 1 | 1 | 1 | 1 | 1 | 1 | 1 |
| Centralization Degree | 0.06927023 | 0.064132654 | 0.1334778 | 0.06181354 | 0.08817454 | 0.095475512 | 0.0791377 | 0.0712395 |
| RM (Relative Modularity) | 5.2693558 | 6.040679146 | 3.17505499 | 4.79215729 | 12.0494888 | 10.78020977 | 8.7625972 | 8.561277694 |

## Conclusion

This study contributes to a deeper understanding of the intricate interactions between bacteria and plants in the context of drought stress. Specifically, our metagenomic analyses in the fields of Louisiana under drought conditions have provided insights into microbial diversity and primary bacterial components of the rhizosphere under such arid circumstances. The findings from our co-occurrence network analyses have corroborated and strengthened our understanding of the microbial dynamics in response to drought stress. Our study establishes a foundation for further research and highlights the importance of defining microbial diversity in the context of drought stress, particularly in regions like Louisiana facing challenges related to drought conditions.

